# Mitochondrial genome evolution in the Diprionidae: Major gene rearrangement in the basal Hymenoptera

**DOI:** 10.1101/2023.03.14.532325

**Authors:** Min Li, Gengyun Niu, Min Xu, Mengxuan Dai, Xinghong Jiang, Yong Ma, Guanliang Meng, Meicai Wei

**Affiliations:** Laboratory of Insect Systematics and Evolutionary Biology, College of Life Sciences, Jiangxi Normal University, Nanchang 330022, China; School of Computer Information Engineering, Jiangxi Normal University, Nanchang 330022, China; Center of Taxonomy and Evolutionary Research, Zoological Research Museum Alexander Koening, D-53112 Bonn, Germany

**Keywords:** gene rearrangement, *nad2*, evolutionary pattern, phylogeny, Ka/Ks

## Abstract

In comparison to other non-parasitic basal lineages, Apocrita have consistently demonstrated a greatly accelerated rate of gene rearrangement. A number of mechanisms or correlates have been proposed for this observation, such as oxidative stress tolerated by exposure to the host immune system might lead to a high proportion of rearranged mt-genomes. Our studies reveal that gene rearrangements involving the protein-coding gene are present in the basal Hymenoptera lineage based on enriched sampling. We speculate the processes of diversification of rearrangements in the vicinity of *nad2* involved tRNAs and NCRs by producing the chronogram of Diprionids. Furthermore, we investigated the relationship between rearranged genes and their nucleotide sequences. In conclusion, we demonstrate the great potential of gene order and associated sequence features as phylogenetic markers in the study of Hymenoptera evolution, offering a new perspective on studying organisms that undergo frequent gene rearrangements.

## 1. Introduction

The studies of diminutive mitochondrial genomes, the fledglings of genomics, have led to many ‘textbook descriptions’ [1]. Rapid advances in sequencing technology have dramatically expanded those findings and deepened or refreshed our understanding of the typical features, leading us to explore the mechanisms or patterns of evolution that are more explanatory. Genomic architecture is probably the most fruitful Genome-level character, with new findings including two co-existing and divergent mt-molecules in single individuals in vertebrate [2] and invertebrates [3], changes in the transcriptional orientations of genes [2], architectural variation of individual genes [4], etc.. Recent meta-analysis [5,6] reinforced more variable rearrangements than anticipated, and identified convergence events. However, as taxonomy experience shows, evolution patterns differ along ranks [7,8]. That means that patterns revealed by meta-analyses of higher taxa, i.e. phyla or orders, can obscure patterns found within order or families. It may thus be possible to obtain new insights into evolution patterns and molecular mechanisms base on dense sampling for representative lineages.

The mt-genome of Hymenoptera is a good model for investigating gene rearrangements [9–12]. It is large enough to accommodate repetitive patterns and complexity, while the basal taxa beyond the massive radiation are small enough to allow for near-complete genus-level sampling in phylogenetic analyses. Recently, a greater than expected diversity of gene rearrangements has been reported in basal Hymenoptera, including inversion, transposition, inverse transposition, shuffling of tRNA clusters, and even inversions of protein-coding genes [13]. This prompts caution regarding the previous conclusion that gene rearrangements occur mainly in parasitic taxa [14]. Inversions of protein-coding genes have been observed in the mt-genome of Diprionidae, which forms a monophyletic group with Cimbicidae and is located at the base of Tenthredinidae.

The internal relationships of Diprionidae have not been fully investigated, but traditionally they have been divided into two subfamilies. Diprioninae are multi-generational over the year, while Monocteninae have only one generation per year. Molecular phylogenetic inferences [15] support a paraphyletic Monocteninae, with the representative of Augomonoctenus being separated at the most basal position of the family. Diprionidae is unique in that all of its species have reverted to a lifestyle of feeding on gymnosperms [16]. Did retreating to less prosperous hosts subject them to resulting selective pressures that prevented them from developing their diversity, and how did this astonishing reversal occur during evolution?

Here we provide a robust mt-genomic dataset including 33 sampling represent 11 genera or same level for inferring the evolution of the Diprionidae and reconstructing gene rearrangement pattern and ancestral gene states in a phylogenetic framework. The gene rearrangement diversification was analysed from multiple perspectives, including nucleotide composition (polymorphism analysis), Ka/Ks calculated by sliding window and rearrangement frequencies. We aim to describe and quantify the diversity of the gene rearrangements by tracing evolutionary history, to detect sequence footprints generated by gene rearrangemtns, and discuss their phylogenetic significance. This will provide new insight for the application of gene rearrangements in the study of the evolutionary history.

## 2. Materials and Methods

### 2.1. Mitogenome sequencing, assembly and annotatio

In total, 33 Diprions were analyzed across two subfamilies and eleven of twelve genera (Table 1). Among them, 13 are newly sequenced and assembled, and 15 are assembled based on raw data from UCEs, transcriptomes, and Hic libraries. Total genomic DNA of a single specimen was used for library preparation with insert size of 250 bp following the manufacturer’s instructions, and then 150 bp PE sequenced on an illumina Nova 6000 platform at Berry Genomics Corporation, Beijing, China. The raw reads were quality checked using FastQC.

**Table 1.**
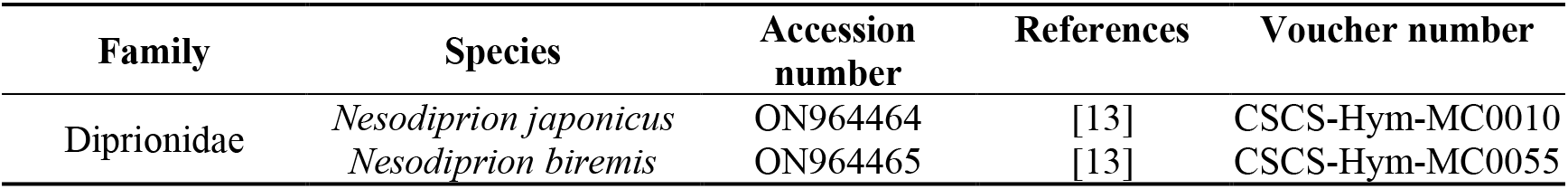
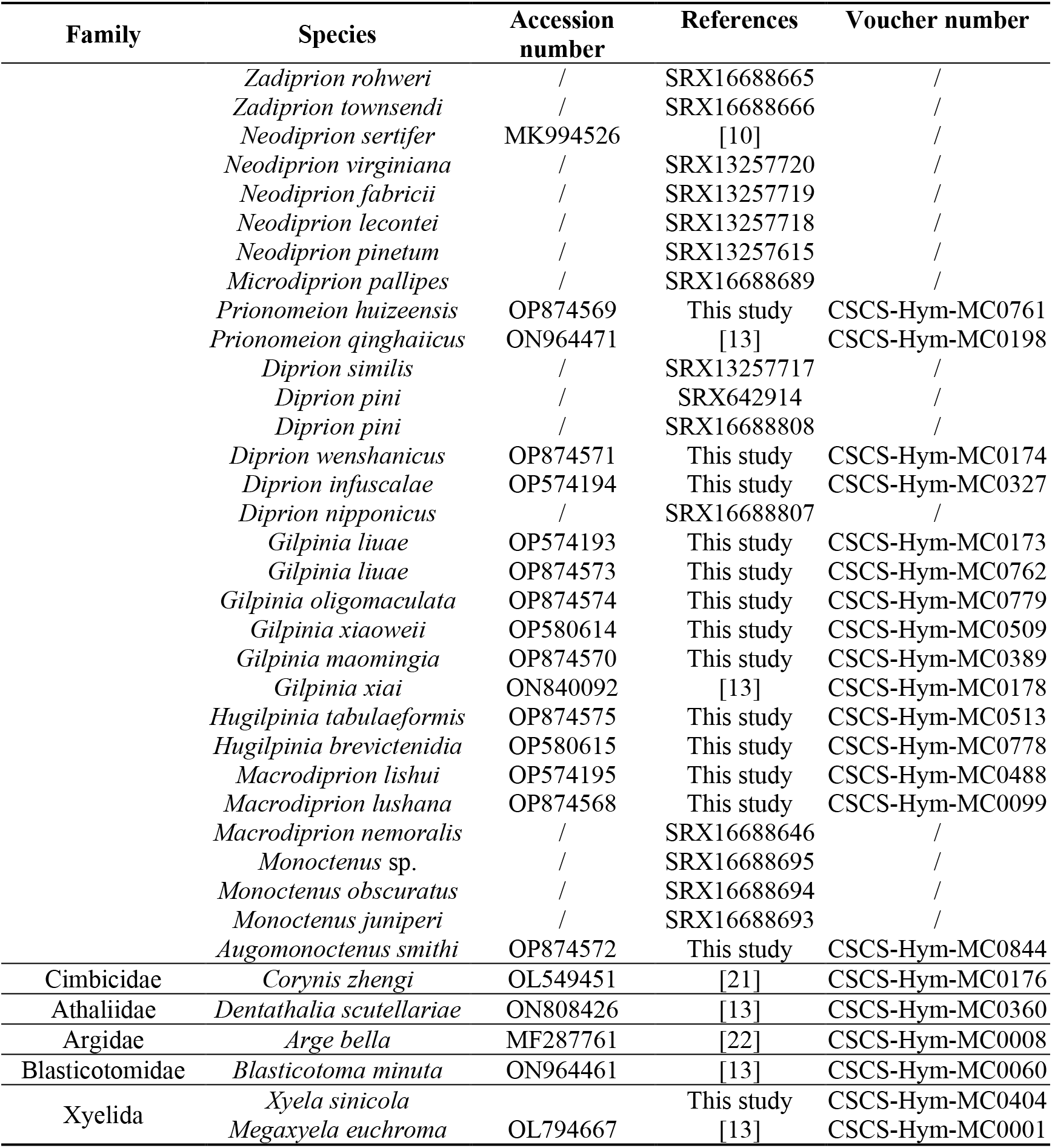
Summary information of mitogenomes used in phylogenetic analyses

Mt-genomes in this study were assembled using two parallel strategies: *de novo* genome assembly by applying the compulsory MitoZ [17] and either NovoPlasty [18] or GetOrganelle [19], and mapping the reads to the reference using Geneious Prime v. 2022.2.1. Secondary verification and assessment of the sequencing coverage were performed for gene rearrangement regions, the absence in *de novo* genome assembly, and other aberrant results. the tRNA reported as absent in this study have been assembled using both the homologous gene and the flanking regions of the closely related species as the reference sequence, and return invalid results. Due to the high uncertainty of CR, the potential for multiple repeats within it was not considered here and only the most parsimonious version was recovered. PCG, tRNA and rRNA were hand annotated according to homology to validate automated annotations in Mitos [20]. For additional copies of tRNA, sequence comparisons with the homologous gene from related species were conducted.

### 2.2. Comparative analysis of mitogenome and visualisation

The sequences were aligned using Geneious with the default parameter, and manual adjustments were made when necessary. Geneious was also used to calculate consensus, identical sites, and pairwise identities.

Using PAML 4.2 [23], we estimated pairwise non-synonymous substitutions per non-synonymous site (dN), synonymous substitutions per synonymous site (dS), and the ratio of these values. For fast base substitution, a KaKs calculator [24] implementation was developed in Python using a sliding window that moves along every pair of sequences. For each window, point marks are used in order to generate a continuous curve. This curve is then combined for all specified samples for comparison.

Protein structures were predicted by SWISS-MODEL [25] and DeepTMHMM [26], using PyMol (*www.schrodinger.com/pymol*) for editing and visualization.

### 2.3. Phylogenetic analysis and ancestral states estimation

DAMBE [27] was used to test for severe substitution saturation in the PCG of mtG. The unsaturated aligned sequences of PCG were concatenated using SequenceMatrix v. 1.7.8 [28] to form a single supermatrix. To improve the accuracy of the phylogenetic analysis, GTR+I+G was selected as the best-fit model using PartitionFinder and subjected to maximum likelihood (ML) analyses and Bayesian inference (BI) analyses using the standard partitioning schemes “BIC” and “AICc”, respectively. ML analysis was performed using the IQ-TREE web server [29], based on a probabilistic modeling approach to calculate the relationships between individual species. Default parameters were used except for a perturbation strength of 0.1 and an IQ-TREE stopping rule of 1,000. BI analyses were performed using MrBayes v. 3.2.2 [30] for Markov chain Monte Carlo (MCMC) simulations based on probabilistic and Bayesian statistical theories. A representative phylogenetic tree was derived by constructing a consensus tree.

We also tried some alternative matrixs to deleted the fragmented sample. The results did not strongly affect the inferences (data not shown); and thus, we only use the result from the matrix 1.We chose to map gene arrangement states onto the tree of matrix, which contains the most OTUs (operational taxonomic units), even if those OTUs from the UCE assembly didn’t always yield a definitive gene arrangement state. On terminals, gene arrangements were counted and grouped based on consistency. Then the gene arrangement states on each node were inferred based on the maximum parsimony. Species divergence times were estimated using the PAML package MCMCTtreeR with three fossil calibration loci, 237~251 Ma, 185~235 Ma and 165~170 Ma, respectively. Evolution times were aliquoted in units of 5 Ma and the gene rearrangement of all nodes in each interval were quantified using qMGR [31].

## 3. Results

### 3.1. Archtecture of mitogenomes in Diprionidae

In this study, we assembled a total of 27 diprionids mt-genomes, using various sequencing approaches. Specifically, nine mt-genomes were obtained from UCEs, one from the transcriptome, five from Hic libraries, and 13 from whole genome sequencing (WGS). Although UCE sequencing may be an efficient method for obtaining mitogenomes, our results demonstrate a low degree of assembly completeness. Only two of the nine samples retrieved all 37 genes, while others had varying degrees of gaps, with the shortest mt-genome containing only nine PCGs. The Hic libraries yielded complete mitogenomes for three Nesodiprions, but severely deficient ones for the remaining two Diprions. On the other hand, WGS produced more complete mitogenomes, with nine resulting in circular complete mitogenomes based on the minimal CR protocol, and four others yielding all 37 genes. The transcriptome data source yielded a poorer assembly, with 11 incomplete PCGs but no tRNAs. Additionally, we observed heteroplasmy in Gilpinia maomingia, with the majority of variation occurring in the PCGs. We retained degenerate bases in those nucleotide positions in order to generalize their variation.

In addition to the previously reported diprionid mt-genomes [10,13], a comparative study of mt-genomes was conducted on 33 diprionid species from eleven genera (Table 1). Overall, diprionid mt-genomes exhibit a high number of non-coding intergenic spacer (IGS) regions, and unlike most other sawflies, the majority of diprionid mt-genomes possess two major non-coding regions (NCR). Multiple gene rearrangements, including a significant reversed *nad2*, have also occurred in numerous species. The length of the Diprionid mt-genome varies considerably, ranging from 15,344 bp (*Gilpinia Xiai*) to 17,991 bp (*Prionomeion qinghaiicus*), primarily due to the presence of multiple IGSs and major NCRs of varying sizes. Several notable features can be observed in these non-coding regions:

1. It is common for Diprionid mt-genome to have two major NCRs between 472bp and 1240bp in length, and NCRs at the same position are highly similar within the genus. Two Nesodiprion species, for example, have NCRs between *trnM* and *trnY* that are 454 and 449 bp, respectively. Their alignment is 456 bp long and contains 389 identical sites. Similarly, the length of the major NCRs in Mt-genomes of Neodiprion ranges from 977 to 1002, with 82.3% of the sites being identical within a 1059 bp alignment.
2. Gilpinia species showed moderate similarity among the major NCRs at the same position, but the two major NCRs themselves were somewhat consistent. The major NCR located between *nad2* and tRNA cluster *MIY* had 529 identical sites relative to the total length of 1162 alignment (49%), whereas the one located between tRNA cluster *MIY* and *cox1* had 891 identical sites relative to the total length of 1358 alignment (65.6%). Yet the similarity between two NCR consensus is 54.2% with 45.3% of the sites being identical.
3. Unlike most sawflies mt-genomes, where the major NCR starts from the 5’ end of rrnS, the Diprionid mt-genome has only short NCRs, with length less than 70 bases, inferred from the secondary structure of *rrnS*, with some exceptions, such as *Neodiprion sertifer* (650 bp) and *Augomonoctenus smithi* Xiao et Wu (472 bp).
4. Genes that have undergone rearrangements in diprionids are surrounded by many IGSs. In the genus Gilpina, for instance, the *trnE* gene has been shifted to the third position from the fifth position in the tRNA cluster between *nad3* and *nad5*, resulting in longer IGSs and a loose arrangement of tRNAs.

Diprionids are characterized by their reversed *nad2* apart from the loosely packed mt-genome. To confirm the reversal, three methods were used: 1. by checking the mapping of reads assembled without reference. 2. by extending both ends by referencing the region between *nad2* and *cox1* and the region between *nad2* and *rrnS*. 3. for some samples, by PCR (unpublished). However, due to the unavailability of raw data, validation for *Neodiprion sertifer* could not be carried out. Generally, in Diprioninae, *trnWC* and *trnIM* as the subsets form a common interval [32] with *nad2*, which undergoes initial complete reversal, followed by various translocations and reversals within or between the small subsets, leading to diverse rearrangements between *rrnS* and *cox1*. Apart from the common interval, other gene rearrangements are also clade-specific.

### 3.2. Diprionid Phylogeny and rare genomic changes as features of higher taxa

Phylogenetic inference of PCGs was performed using 39 genomes, sampling 6 outgroups and 33 Diprionids taxa (Matrix 1, 11622 bp). The initial set was filtered into WGS source data for 27 Diprionidae taxa (Matrix 2, 11607 bp), and only the Diprioninae were retained (Matrix 3, 11403 bp). We utilized the infinite mixture site heterogeneous model (CAT-GTR) for all matrices, while ML analysis and BI analysis were used for the latter two matrices, respectively.

The monophyly of Diprioninae, Monoctenus, and Augomonoctenus was consistently recovered in all five analyses. Augomonoctenus was found to be the basal clade of the family, while Monoctenus was the sister group to Diprioninae (bootstrap frequency [BS] > 95%; posterior probabilit > 0.95). The relationships within Diprioninae varied across topologies, but the monophyly of each genus, and three clades was consistently confirmed: 1. Nesodiprion and Neodiprion, 2. Prionomeion and Diprion, 3. Macrodiprion (Hugilpinia + Gilpinia). Clade 1 and 2 were sister groups, while clade 3 was basal, which was supproted by all anaylises except for the ML inference for matrix 2. Whith dense sampling in matrix 1, clade 1 was turns to ((Zadiprion + Neodiprion) Microdiprion) Nesodiprion (Figure S1). In order to assess the topologies, we refined several key morphological features. Based on their distribution along the topology, we concluded that the bayesian inference of matrix 3 under the heterogeneity model was the most reliable, which was then further analyzed.

On the phylogenetic tree, gene rearrangement shows step-like changes, resulting in each clade having its own characteristic pattern (Fig. 1). All Diprionids had their *trn<u>P</u>T* swapped relative to the Hymenoptera ancestral types. After that, *trnAR* was switched in all clades except Augomonocenus, which is the most basal. Once the sub-basal Monoctenus was removed from consideration, a ground pattern of reversal was observed in Diprioninae, with *trnQ<u>W</u>C* located upstream of the reversal *nad2*, and *trn<u>IMY</u>* downstream. On this basis, *trnC* translocated to the upstream of *trnQ* in Zadiprion in Clade 1. With the exception of incomplete genomes (*Zadiprion rohweri* and *Microdiprion pallipes*), the major NCR of all other genomes lies between *trn<u>M</u>* and *trn<u>Y</u>*.

**Fig. 1.**
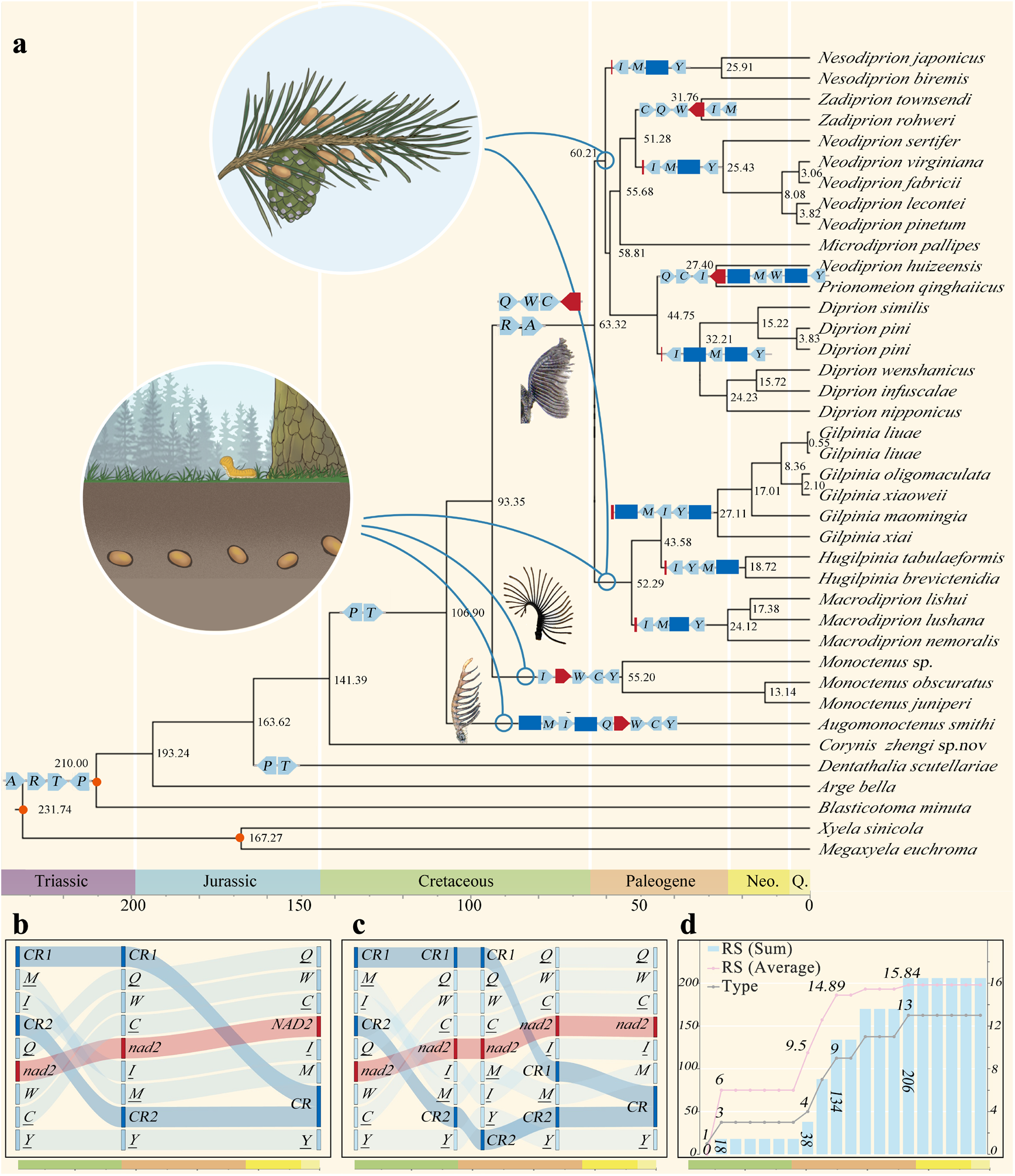
Timescale of Diprionidae evolution and intensity of gene rearrangement. (a) Time-calibrated phylogeny of Diprionidae is based on the Bayesian tree infered with matrix 1 and divergence times are estimated with MCMCTree. Placements of fossil calibrations are indicated by red spots and gene rearrangement of each clade are showed on the respective nodes; Two different locations of cocoon in Diprionidae are marked on the tree. (b) The predicted gene sequences of Nesodiprion and Neodiprion changed over time. (c) The predicted gene sequences of Diprion changed over time. (d) The observed gene rearrangements in Diprionidae are counted using qMGR based on ancestral gene patterns and counted every five million years from the expected time of first appearance of Diprionidae.

Clade 2 has undergone a striking rearrangement around the *nad2* gene, with *trn<u>W</u>* relocating from the *trn Q<u>W</u>C* cluster to the downstream of *nad2*, setting in the middle of *trn<u>M</u>* and *trn<u>Y</u>*, Meanwhile, *trn<u>I</u>*, originally flanking *nad2* at the downstream side, has moved upstream and inserted into the middle of *trnC* and *nad2*. In *Prionomeion qinghaiicus, trn<u>M</u>* and *trn<u>W</u>* have shifted even further upstream, forming a cluster of five trns positioned upstream of *nad2*. Both Prionomeion genomes possess two major NCRs, with the 700bp NCR located upstream of the longer NCR exceeding 1000 bp. These NCRs are located upstream and downstream of *trnI* in *Prionomeion qinghaiicus* and of *trn<u>M</u>-trn<u>W</u>* in *Prionomeion huizeensis*, representitvely.

Clade 3 exhibits increased flexibility in the relative positions within *trn<u>IMY</u>* cluster. Hugilpina mtgenomes possess only one major NCR located downstream of the *trn<u>IYM</u>*, whereas Gilpinia mtgenomes have two major NCRs flanking the *trn<u>MIY</u>*. In addition, the major NCRs of the basal Macrodiprion are situated within the *trn <u>IMY</u>* cluster.

Aside from the synapomorphic rearrangements around *nad2* observed in each genus, the tRNA cluster between *nad3* and *nad5* is distinctive in Gilpinia, with *trnE* relocating from its ancestral position (between *trnS1* and *trn<u>F</u>*) to a derived position (between *trnA* and *trnN*).

### 3.3. Comparative sequence features in the mitogenomes in Diprionidae

Using *Dentathalia scutellariae* as a reference, we calculated Ka/Ks ratios for Diprionidae. Across all 13 protein-coding genes (PCGs), the extreme values differed by 0.922, from 1.118 (*atp8*) to 0.195 (*cox1*). We performed the same calculations for Cimbicidae, the sister group of Diprionidae, and the differences between the genes of the two populations showed approximately the same trend.. (see Figure S2). When calculated pairwise for each species, the values also remained relatively low. Even for *nad2*, which had the high ratio, the comparison ranged from 0.536 (*Prionomeion huizeensis*) to 0.960 (*Diprion nipponicus*), with a mean of 0.754 (Table S1). The Ka/Ks ratios of *atp8, nad6, nad2* and *nad4l* ranked in the top four.These findings indicate that purifying selection has occurred, both for individual genes and individual species.

However, when we applied a sliding window to perform higher resolution statistics on the sequences, we observed that the sub-regions with Ka/Ks values above 1 emerged (Fig. 2). This observation suggests that although the entire genes followed neutral evolution, specific regions underwent amino acid fixation during evolution, indicating that there is variability in the selection to which different structural and functional protein domains were subjected. Specifically, *nad2* exhibited a clade-specific variations throughout the family, three to five waves appear in the Ka/Ks>1 section, and have a maximun peak at the end of the sequence.

**Fig. 2.**
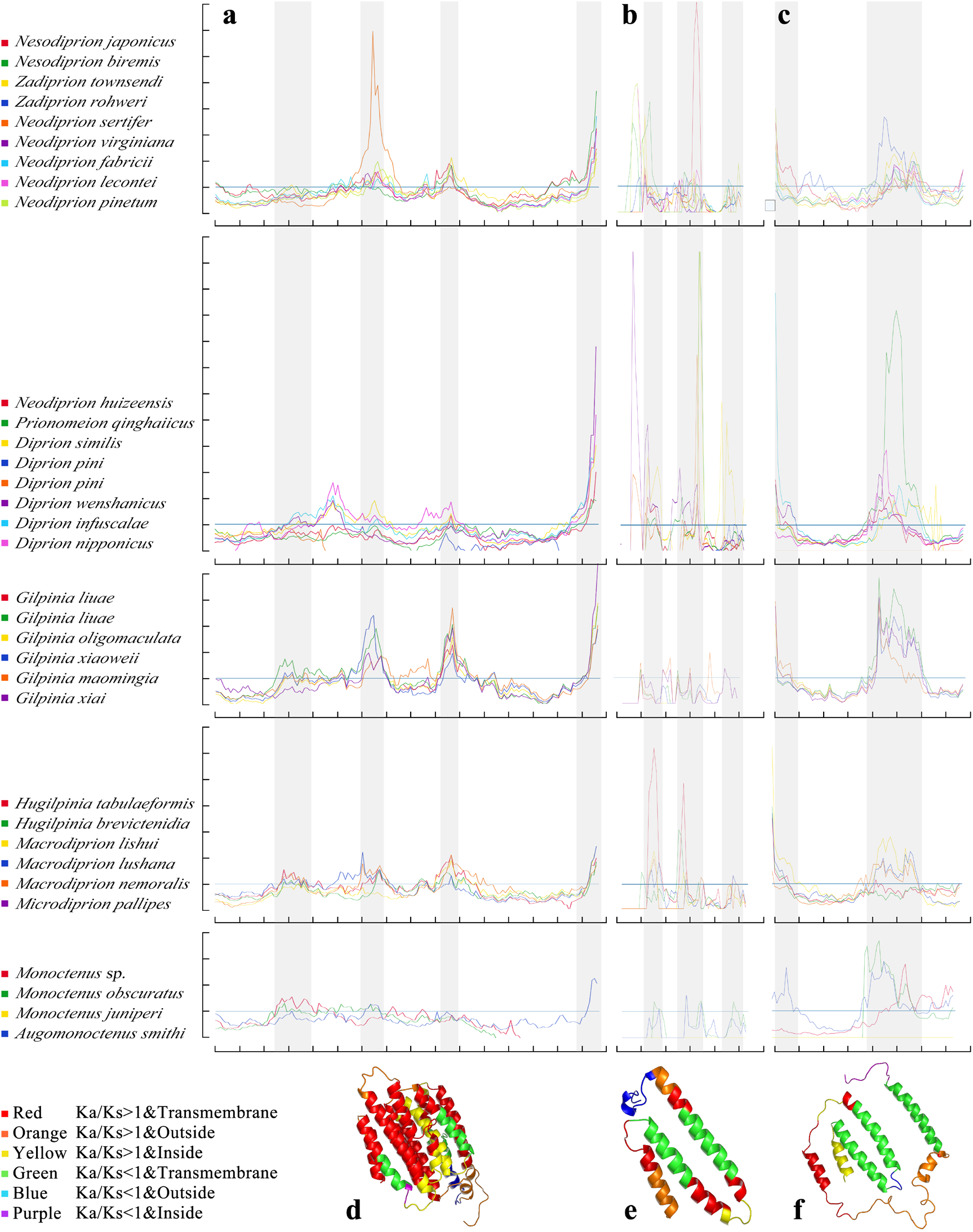
Sliding window analyses of Ka/Ks and 3D models in Diprionids. Sliding window analyses of Ka/Ks ratios among (a) *nad2*, (b) *nad4l* and (c) *nad6* of 33 species from Diprionidae, using *Dentathalia scutellariae* as reference genome. The window size used are 132,45 and 90, respectively. Species that displayed similar curves are grouped in a clade. The Y-axis showed the Ka/Ks ratio with each scale representing 1 and the x-axis showed the nucleotide sites at which the alignment sequence begins with each scale representing 10. Protein coding genes (d) *nad2*, (e) *nad4l* and (f) *nad6* of *Nesodiprion biremis* are predicted by using SWISS-MODEL. The topological structure and Ka/KS values were color-coded by Pymol according to the results of DeepTMHMM.

This pattern shows similarity in related taxa and we divied all the species involved in the calculation into five clades. For example, Diprion and Gilpinia both displaced four waves in similar intervals, while Monoctenus and Neodiprion+Neodiprion have one less in relative terms, located in intervals 20-40 and 87-105, respectively. Similar phenomena were also observed in *nad4l* and *nad6*, where waveform similarities were observed within clades, but variability was seen between clades. This repeatable variability was hardly observed in other PCGs, like *cox1*, which repeats the similar waveform patterns across the entire family.

Considering that the conserved nature of reproducible differences appears in genes with positional correlation, we therefore predicted the structure of the proteins encoded by these genes and also tested for the presence of co-expression between these genes. to identify more precisely whether there is co-evolution in the above sub-regions (Fig. 2). The results show that *nad2* and *nad4l* and *nad6* are closely related and overlap each other, which may indicate that they are functionally and structurally interdependent and adaptive. In the case of *nad2*, most of the regions with Ka/Ks values higher than 1 are located in the transmembrane region.

## 4. Discussion

### 4.1. Phylogenetic Framework of Diprionidae

Through proportional sampling of lineages within the clade, we were able to cover 11 of 12 genera or equivalent rank. All analyses produced a consistent framework, indicating that Diprionidae is composed of a diverse, monophyletic Diprioninae (including 8 genera and over 140 species) and sequentially, two poorly diversified basal clades (comprising 3 genera and 16 species). These results are consistent with previous finding based on short fragments of mitochondrial and nuclear genes [15]. As such, the taxonomic system that divides Diprionidae into two subfamilies was rejected, as the subfamily Monocteninae was not found to be monophyletic.

The observed pattern, characterized by a sparse genealogy of multiple ancient species at the base and a diversified clade, is supported by a suite of morphological, biological and genomic evidence. Morphologically, each of the three evolutionary clades possesses unique male antenna type (Fig. 3), a key diagnostic feture for higher taxa in the basal Hymenopteran family and subfamilies. Furthermore, three additional plesiomorphic characters are present in the basal two clades are: an ancient pattern of mesoscutellum and metascutellum, a smooth and hardly punctured body surface, and a simple penis valve. Conversely, in Diprioninae, the mesoscutellum and metascutellum are distinctly modified, the body surface is densely and strongly punctate, and the penis valve are specialized. Both basal clades have typically one generation per year, with *Augomonoctenus smith* requiring up to two years to complete its life history [33], and cocooning 5 to 10 cm below the soil surface. Augomonoctenus larvae bore cones of *Cupressus* [34] or *Libocedrus* [35], while Monoctenus larvae feed on *Juniperus* needles [36]. The diverse Diprioninae are mostly multi-generational, with Nesodiprion even having 5-6 generations per year [37]. Diprioninae larvae are needle-feeders and no longer cocoon in the soil, but instead have shifted to the ground [38], litter [39,40], or even needles [41].

**Fig. 3.**
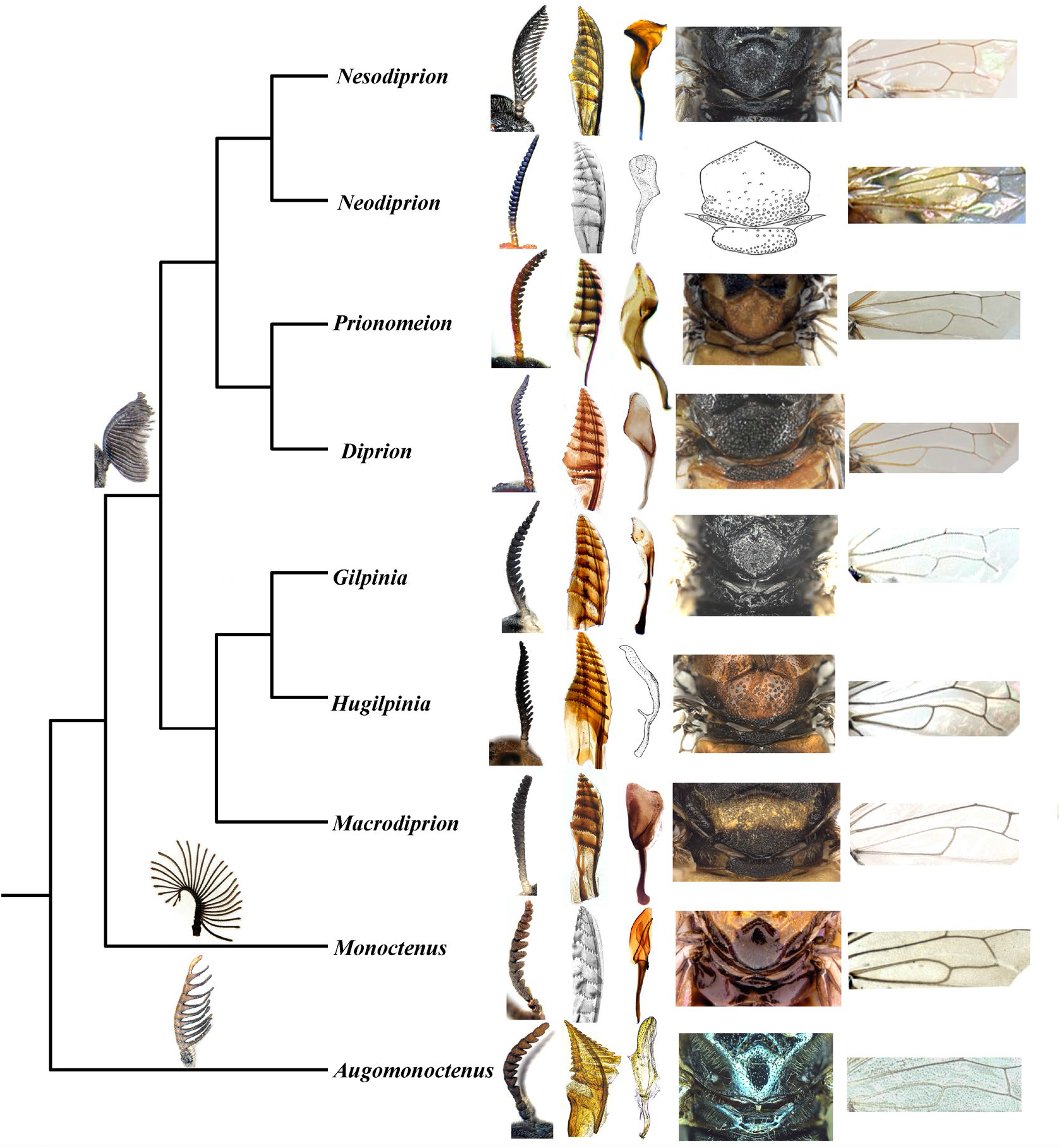
The morphological characteristics of the Diprionidae: male antennae, penile valve and vein.

In terms of gene order, the dense taxonomic sampling has filled in the gaps in the otherwise blurred evolutionary history. Table S1 detailed the inferred mtgenomic organizations from nearly complete or complete sequences of 33 Diprionids. However, the UCE and Hic rawdata failed to obtained some sequences, particularly those near *nad2*. While the impace is perhaps minor in genera such as Diprion and Neodiprion, where no significant difference was found when comparing mtgenomes with the genus. Gene arrangements may yet be underestimated in Microdiprion and Zadiprion. Moreover, caution should be exercised with genera that have small sample size, such as Prionomeion, which displayed gene arrangements divergence in only two samples. At least eleven types of gene arrangement have been observed, with each genus having at least one exclusive rearrangement pattern. The two basal clades retained the basic pattern, while the entire Diprioninae share the derived inversion of *nad2*. Furthermore, additional rearrangements involing tRNA and NCRs led to different arrangement patterns among lineages. Some genera also exhibit characteristics that are distinctive from the general state. The gene rearrangement in the Diprionidae family has several notable features, including: 1. A derived *trnPT* is shared by the family; 2. Two major NCRs are commonly found, including the two basal lineages; 3. Diprioninae share a derived *trnRA*; 4. *trnE* is translocated within the cluster in the Gilpinia.

### 4.2. Pattern and evolution pathway of gene rearrangements in Diprionids

The family Diprionidae provides a good model system for studying gene rearrangements, as it includes: 1. a lineage containing a high frequency of gene rearrangements and other lineages that maintain their ancestral status. 2. intermediate types of progressive rearrangements can be observed within the lineage of high frequency gene rearrangements. 3. rearrangement types are conserved within evolutionary branches. This model enables us to quantify the frequency and intensity of rearrangements over time and infer the basic patterns (Fig. 1).

Our results confirmed previously identified hot spots, such as the most mobile IQM tRNA gene cluster, and the *nad3–nad5* junction [42], but not *trnK* and *trnD* [9]. However, shared gene order within the hot spot may result from different evolutionary pathways. For instance, the *trn<u>I</u>-trn<u>M</u>*-NCR-*trn<u>Y</u>* is shared by Nesodiprion, Neodiprion and Macrodiprion. By tracing the path, we found that if the organization of Augomonoctenus is assumed to be the ancestral type of the family, the former two genura underwent only one rearrangement event no later than 63 Ma ago, while the latter underwent at least three changes at varying intervals. It becomes more complex if we consider the NCR. Although NCR are both located between *trnM* and *trnY*, their lengths differ. In Nesodiprion, they are approximately 450 bp, whereas in Neodiprion, they are around 1000 bp, which is twice as long as in Nesodiprion. There appears to be no evidence to suggest that the longer NCR originated from the loss of another ancestral NCR or from the merger of two NCRs.

Observation of gene rearrangements over time reveals that in the Cretaceous, they occurred only at the crown node of the family and then remained silent until the Paleogene, when their frequency increases linearly increased with time. As shown in the Fig. 1, total rearrangement scores (RS) in blue and type statistics in orange showed at least two brief pauses, while average RS scores in purple showed marginal effects soon after the first contemporaneous pause.

The latest rearrangement occurred at 27 Ma and extant species diversity is mainly derived from Neogene and Quaternary from 25 Ma. Therefore, extant species richness not correlated with rearrangements, suggesting that gene rearrangements are a feature of evolutionary branches and do not carry relevant signals for species differentiation within them.

The stepwise scenario prompts a reassessment of convergence [43], previously understood as the convergence of similar rearrangements occurring simultaneously in distant taxa [6] and assigned a phylogenetic significance weighting [44]. We have, however, discovered that rearrangements are frequent [45] and favor hotspots [9], but are random in lineage [46]; if this bias is attributed to molecular mechanisms, the adaptive value of the new orgnazition may depend on specific molecular features, rather than responses to ecological challenges [47]. Unlike the generally mt genomes with tightly arranged gene, Diprionidae has multiple intergenic sequences. The potential homology between IGS and major NCRs suggest that these intergenic regions play a crucial role in regulating the expression of nearby genes [48]. Based on the similarity of major NCR and tRNAs, we can speculate that pseudogenes were once present, and these intervals then shrank over time under deletion pressure to maintain genome compactness [49]. Furthermore, when considering IGS and major NCR, similar arrangements exhibiat different degrees of polymorphism. Therefore, assuming those mobile elements underwent neutral evolution and deleting them to reduce noise [6,50] may thus lead to the lose valuable phylogenetic signals or miss relics of the rearrangement event.

Gene rearrangements are rare [51], making them a useful tool for establishing a link between ‘adaptive radiation’ and its ‘key innovations’ [52]. Diprionidae, as a model, offers a perspective that underscores the necessity of defining the the unit of evolution beforehand. While gene rearrangements appear to be infrequent in genera, they are widely observed at higher taxonomic ranks. At the family level, the intensity and frequency of gene rearrangements linearly increases over time with non-episodic pauses (Fig. 1). This saltatory (non-clocklike) manner [53] has also been observed in nematodes [54], snails [55], and Salamanders [56], indicating that gene rearrangement is temporally ans spatially repeatable. In other words, ecological factors, adaptive landscapes, or evolutionary opportunities may shape similar patterns of gene arrangement.

### 4.3. Sequence footprints and application of gene arrangments in phylogenetic inference

It was believed that all the variations observed in the mitochondrial genome had to be neutral due to the fatal impact pf mutations on functions [57]. As a result, mitochondrial genes were thought to be quantifiable, enabling to estimate the timing of species divergence. In contrast, mitochondrial genes typically perform poorly in genomic analyses aimed at identifying the genetic basis of adaptive changes. Our findings demonstrate that although most PCGs underwent purifying selection, certain regions underwent significant positive selection. Furthermore, the pattern of mutation diverged among different clades (Fig. 1). The sequence footprints left behind during rearrangement events may offer more detailed and reliable evidence for inferring the history of rearrangements and the corresponding mechanisms [58]. Divergence occurs in *nad2, nad4l* and *nad6*, suggesting that different genes, or individual loci within genes, underwent varis evolutionary pathways, resulting in inconsistent sequence footprints. This phenomenon essentially rejects the homogeneity of similar rearrangement types. It also suggests the possibility that mitochondrial gene rearrangements occur following a series of changes with buffering effects [59]. Rearrangements and footprints may provide an opportunity to understand the nature of selection operating on mitochondria.

It should be noted that all three genes encode subunits of the respiratory china complex I. however, it is still too early to determine whether there is co-evolution between them.

### 4.4. Evidence of recombination and rethinking TDRL

Low-resolution taxonomic sampling often leads to an overestimation of the frequency of gene shuffling, which subsequently favors the use of the Tandem Duplication and Random Loss (TDRL) model [60–62] to interpret discontinuous evolutiong However, increasing the sample density may provide stepwise evolutionary pathways by filling in gaps. In Diprionidae, rearranged genes can be grouped into several common intervals, while the major non-coding region appears to contain pseudogenes. These observations support the TDRL model, except for the inversion of *trnQ*, which suggests recombination [43,63,64]. Furthermore, multiple hypotheses, such as the Tandem Duplication and Non-Random Loss (TDNL) model [48] and the Dimer-Mitogenome and Non-Random Loss (DMNR) model [49], have been proposed for specific taxa.

Taking a broader perspective of the major eukaryotic groups [65], it is evident that plant mt-genomes, particularly those in vascular plants are frequently rearranged [66–68]. As are those of protists, which also exhibit variability in gene order [69]. Various rearrangements have also been reported in animal mitochondrial genomes, particularly in birds [70], molluscs [1,71] and Insects [45]. Fungi, also are known for their high level of variability [72,73]. And in terms of the mechanism of rearrangement may be explained by the combined effects of non-homologous recombination [74], cumulative duplication [75] (especially in intergenic regions), mobile element dynamics [66], and possibly mitochondrial nuclear gene interactions [76].

Non-adaptive mechanisms may contribute to our understanding of complex gene rearrangement patterns. Although recombination events may be detrimental, only a few of them are adaptive. The discontinuous gene order observed in these events results from both diversification and extinction. Microevolutionary mechanisms may explain diversification, but not extinction, which may explain the lack of direct correlation between rearrangement patterns and biological features. As previously mentioned, a clear correlation was not observed between the number of generations, gene rearrangement, and corresponding sequence feature changes in Diprionidae. Instead, adaptive features such as life history and cocoon location in Diprionidae may have a co-innovative relationship with the genome, rather than a one-way causal relationship [77]. Given that Diprionidae hosts have shifted to gymnosperms, mutual adaptation between them may be explained by a coevolutionary arms race model, where some recombinant mtDNA may have gained replicative advantages. This could lead to their overexpression in early heteroplasmies and an increased probability of fixation as the lineage becomes more homogeneous.

## 5. Conclusions

This study represents the first comprehensive investigation of the mitochondrial genome across a wide range of Diprionid taxa. The unexpectedly discovery of extensive and high-frequency gene rearrangements in Diprionids provides insights beyond the sequence-based approaches for understanding the organisation, genetics and evolution of animal mitochondrial genomes.

Our investigation highlights the derived *nad2* inversion, clade-specific tRNA rearrangements, and repeatable Ka/Ks variation in complex I, all of which provide opportunities for further exploration of the complex relationship between host transfer phenotypes and molecular evolution in Diprionidae.

By examining the changes in gene order at the generic level within a phylogenetic framework and inferring the frequency and timing of gene rearrangements, we propose that genus-level and species-level have distinct evolutionary patterns, but the effects of radiation and natural selection on gene order remain unclear.

## Supporting information

Supplementary Figure 1, Phylogenetic trees of Diprionidae.

Supplementary Figure 2, Ka/Ks average in Diprionidae and Cimbicidae

Supplementary Table 1, Mitochondrial gene order of 33 species from Diprionidae

Supplementary Table 2, Ka/Ks values in Diprionidae and Cimbicidae

## Author Contributions

Conceptualization, M.W.; methodology, G.N. and G.M.; software, M.D., X.J. and M.L.; validation, G.N. and Y.M.; formal analysis, M.L. and M.X.; investigation, G.N. and M.L.; resources, M.W.; data curation, G.N. and M.L; writing—original draft, G.N. and M.L.; writing—review and editing, M.W.; visualization, M.L.; supervision, M.W.; project administration, M.W.; funding acquisition,G.N. and M.W. All authors have read and agreed to the published version of the manuscript.

## Data Availability Statement

The data presented in this study are openly available in Figshare at https://figshare.com/account/home#/projects/149867

## Acknowledgments

The members of the Lab of Insect Systematics and Evolutionary Biology (LISEB) at Jiangxi Normal University are thanked for their contributions to laboratory work. The authors thank Wang Jin for the assistance with figure editing. We thank the anonymous reviewers for their careful reading and many constructive comments.

