## Supplementary figures and images for "Mitochondrial genome evolution in the Diprionidae: Major gene rearrangement in the basal Hymenoptera"

### Supplementary Figure 1, Phylogenetic trees of Diprionidae.

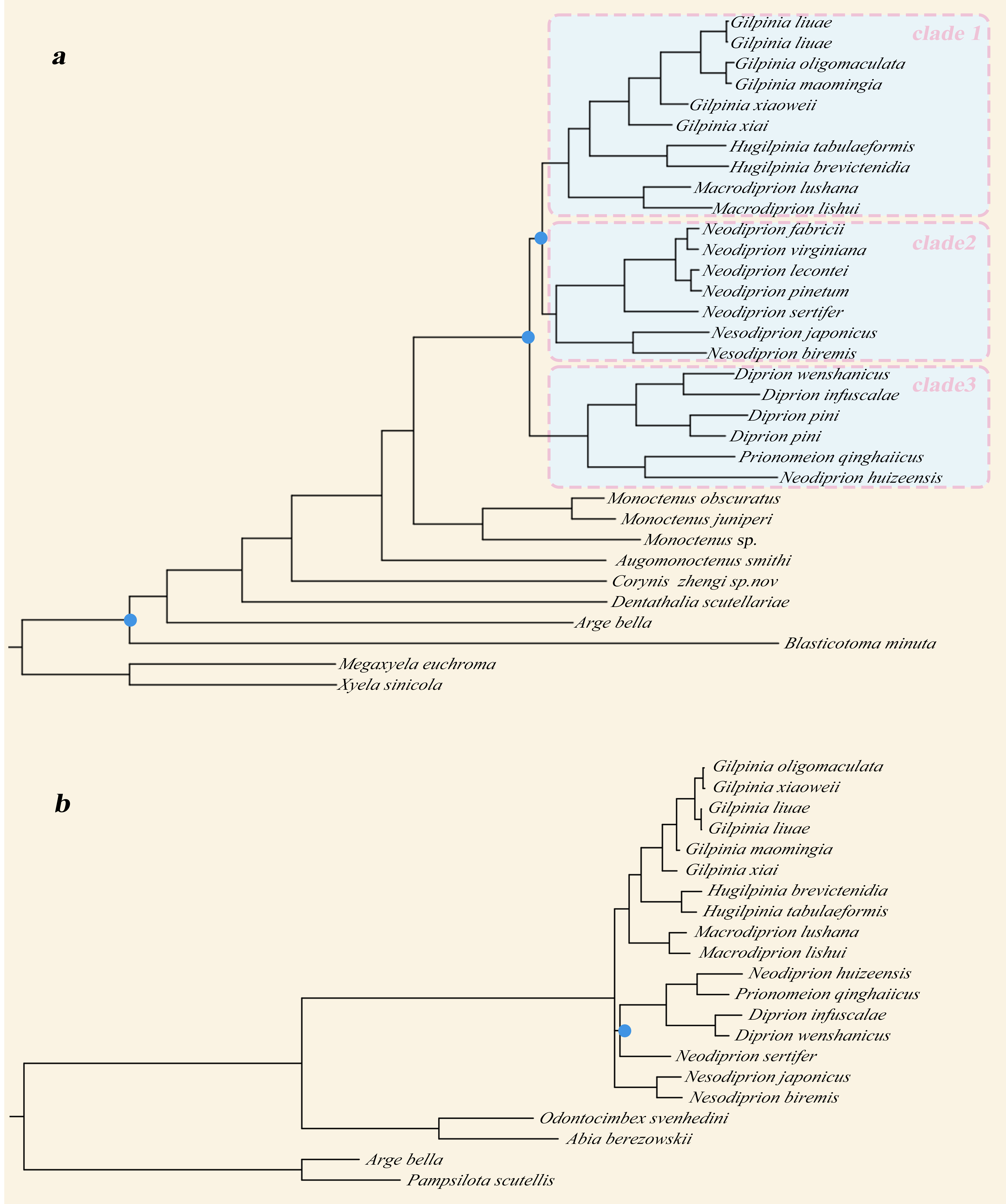

### Supplementary Figure 2, Ka/Ks average in Diprionidae and Cimbicidae

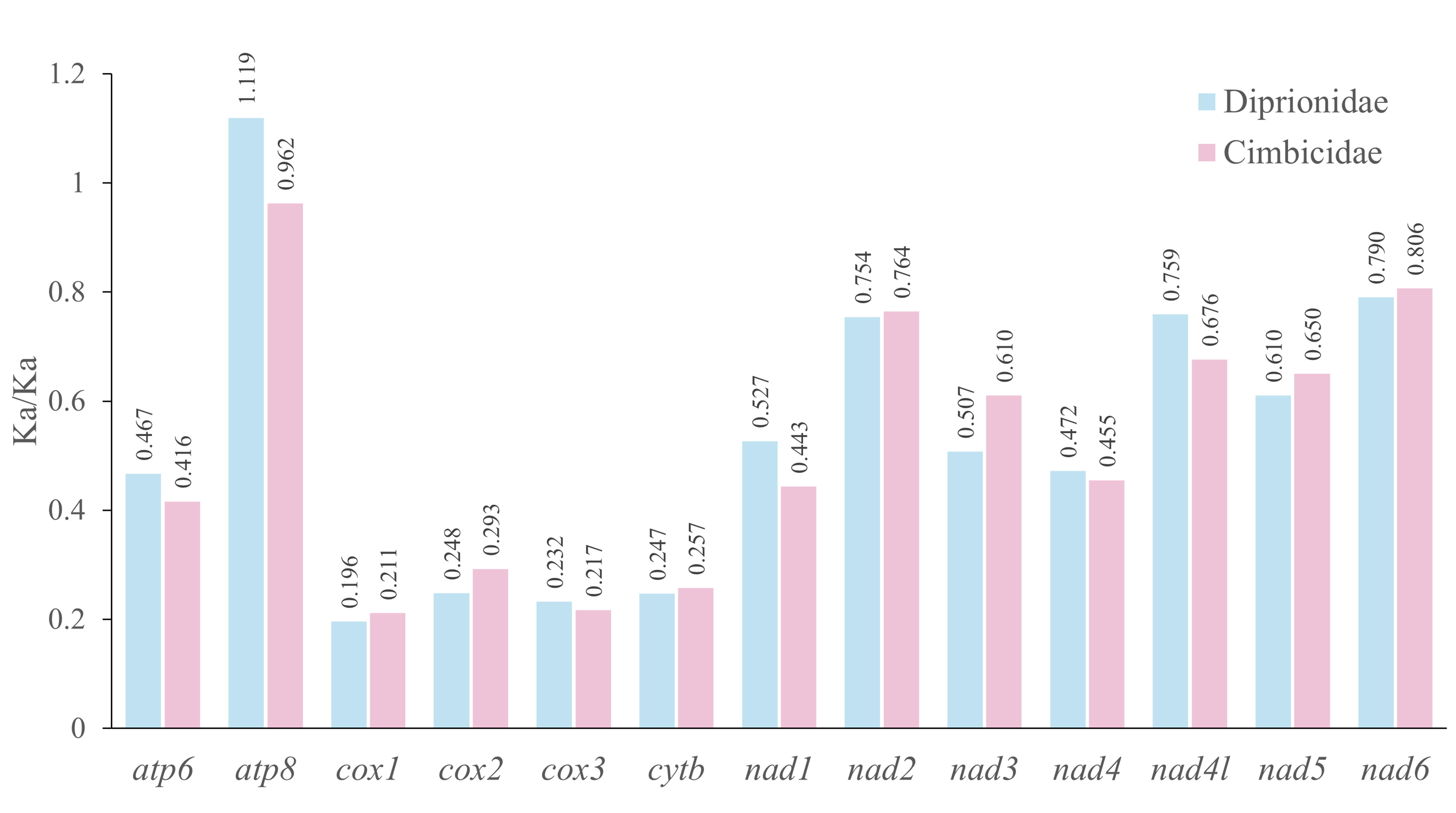
